# Metal contamination and oxidative stress biomarkers in estuarine fish following a mine tailing disaster

**DOI:** 10.1101/2020.06.29.177253

**Authors:** Fabrício Ângelo Gabriel, Rachel Ann Hauser-Davis, Lorena Oliveira Souza Soares, Ana Carolina de Azevedo Mazzuco, Rafael Christian Chávez Rocha, Tatiana Dillenburg Saint’Pierre, Enrico Mendes Saggioro, Fábio Veríssimo Correia, Tiago Osório Ferreira, Angelo Fraga Bernardino

**Author notes:** Corresponding Author: Fabrício Gabriel^1^, Av. Fernando Ferrari, 514, Goiabeiras. Vitória, Espírito Santo, 29075-910, Brasil.

## Abstract

The Rio Doce estuary in Brazil was impacted by the deposition of mine tailings caused by the collapse of a mining dam in 2015. Since the disaster, the estuary is experiencing chronic trace metal contamination effects, but potential trace metal accumulation in fishes has not been reported. Trace metals in aquatic ecosystems pose severe threats to the aquatic biota, so we hypothesized that the accumulation of trace metals in estuarine sediments nearly two years after the disaster would cause contaminant bioaccumulation, resulting in the biosynthesis of metal-responsive proteins in fishes. We determined trace metal concentrations in sediment samples, metal concentrations, and quantified stress protein concentrations in the liver and muscle tissue of five different fish species in the estuary. Our results revealed high concentrations of trace metals in estuarine sediments when compared to published baseline values for this estuary. The demersal fish species *Cathorops spixii* and *Genidens genidens* had the highest Hg, As, Se, Cr, and Mn concentrations in both hepatic and muscle tissues. Metal bioaccumulation in fish was statistically correlated with the biosynthesis of metallothionein and reduced glutathione in both fish liver and muscle tissue. The trace metals detected in fish tissues resemble those in the contaminated sediments present at the estuary at the time of this study and were also significantly correlated to protein levels. Trace metals in fish muscle were above the maximum permissible limits for human consumption, suggesting potential human health risks that require further determination. Our study supports the high biogeochemical mobility of trace metals between contaminated sediments and local biota in estuarine ecosystems.

## Introduction

Estuaries are among the most threatened coastal ecosystems and are continually impacted by anthropogenic actions which often increase the input of organic and inorganic pollutants to the water and sediment (Muniz et al., 2006; Hadlich et al., 2018; Lu et al., 2018). Pollutants released into estuarine ecosystems include trace metals and metalloids that are stable, toxic and persistent environmental contaminants, i.e. they are not degraded, destroyed or subject to transformation (Gómez-Parra et al., 2000; Garcia-Ordiales et al., 2018). The released contaminants typically decrease water and sediment quality, with impacts to estuarine biodiversity and productivity (Lotze et al., 2006). Therefore, understanding the fate and ecological risks of metallic pollutants is critical to access environmental risks associated with their presence in coastal and marine ecosystems (Mayer-Pinto, Matias & Coleman, 2016).

In November 2015, the Rio Doce estuary in SE Brazil was severely impacted by the collapse of a mine tailing dam located 600 km upstream. It is estimated that almost 43 million cubic meters of mine tailings were released, polluting riverine and riparian ecosystems along its descent to the estuary (Do Carmo et al., 2017). The arrival of tailings in the Rio Doce estuary caused immediate impacts to biodiversity and a rapid (<2 days) accumulation of Fe and associated trace metals (Cr, Pb, Mn, and Al) in estuarine sediments (Gomes et al., 2017; Queiroz et al., 2018; Andrades et al., 2020). Nearly two years (2017) after the initial impact, there is evidence for continued chronic effects from trace metal contamination on benthic assemblages (Bernardino et al., 2019), but the extent of the metal contamination and sublethal effects to fisheries in the estuary remain to be determined.

Metals may bioaccumulate in aquatic organisms causing a range of sub-lethal effects. These effects include metabolism depression, diseases and genotoxic damage, and as a result fishes are important bioindicators of metal contamination (Carrola et al., 2014; Lavradas et al. 2014; Ahmed et al., 2015; Gusso-Choueri et al., 2016; Gu et al., 2018). Contamination effects can be immediately detected by sensitive biochemical indicators that are efficient applied to environmental monitoring programs (Van Der Oost, Beyer & Vermeulen, 2003; Hauser-Davis, Campos & Ziolli, 2012). In aquatic ecosystems, fish perform detoxification functions and, when not able, display induced oxidative stress proteins once exposed to metal contamination (Atli & Canli 2008). For example, metallothioneins are low molecular weight proteins that act in the homeostasis of essential trace elements (e.g. Cu and Zn) and in the detoxification of toxic elements (e.g. As, Cd, Pb, Hg, among others; Hauser-Davis et al., 2014; Kehrig et al., 2016). Metallothionein expression increases above certain metal threshold conditions, where the presence of the thiol groups of cysteine residues allows for these metalloproteins to bind to specific metals, protecting the body from metal toxicity through immobilization, metabolic unavailability, and subsequent detoxification, mainly in the liver or organs with equivalent function (Kehrig et al., 2016; Okay et al., 2016; Pacheco et al., 2017; Van Ael, Blust & Bervoets, 2017). Another biomarker, the tripeptide reduced glutathione (GSH, γ-L-glutamyl-L-cysteinyl-glycine), is an important intracellular antioxidant and defense mechanism which intervenes against intracellular oxidative stress-induced toxicity (Lavradas et al., 2014; Kehrig et al., 2016). The sulfhydryl group (−SH) present in cysteine is involved in protective glutathione functions (reduction and conjugation reactions), which provide the means for the removal of many xenobiotic compounds (Meister, 1992). As a result, biochemical biomarkers have become effective tools for assessing toxic effects of metals in aquatic organisms (e.g. Campbell et al., 2008; Lavradas et al., 2014; Hauser-Davis et al., 2016; Okay et al., 2016; Pacheco et al., 2017).

Considering that fish protein is an important source of nutrition for human coastal communities, the evaluation and determination of metal concentrations and their distribution in aquatic fauna are essential for food quality assessments, predictions regarding the toxic potential of these food items for local communities and guide immediate public health actions (Hauser-Davis et al., 2016; Van Ael, Blust & Bervoets, 2017; Coimbra et al., 2018).

Considering the potential long-term effects of trace metals to fisheries, this study quantified metal contamination and the expression of two detoxification proteins on fish from the Rio Doce estuary nearly two years after tailings contamination. Our protocol included (i) the quantification of metal contents in muscle and liver tissue; (ii) the determination of oxidative defense and detoxification biomarkers in the muscle and liver tissue of five fish species consumed by local villagers, and (iii) statistical correlations between tissue metal accumulation and protein expression. Our hypothesis was that chronic exposure to contaminated sediments, 1.7 years after the disaster, would lead to the assimilation of trace metals and expression of oxidative defenses in fish liver and muscle. Additionally, we compared metal and metalloid concentrations of fish to reference values in Brazilian and international guidelines. This study provides a timely and critical assessment of the sublethal impacts and bioaccumulation of metals in fish that are used for the subsistence of villagers that rely on fisheries from the estuary.

## Materials & Methods

### Study area and sampling

The Rio Doce estuary is located in the Eastern Brazil Marine Ecoregion (19° 38’ to 19° 45’ S and 39° 45’ to 39° 55’W). The area has two well-defined seasons, a dry winter (April to September) and rainy summer (October to March), with an average monthly rainfall of 145 mm and average temperature ranging from 24 to 26 °C (Bernardino et al., 2015). The estuary is characterized by a main channel with sand pockets that form at low tide, with salinities typically ranging from 5 to 0 (Gomes et al., 2017). The estuary is used by local villagers as a source of subsistence through fisheries and tourism, and estuarine fishes are an important source of protein (*Mugil* sp. and *Eugerres brasilianus*; Pinheiro & Joyeux, 2007).

The nature of the disaster made the establishment of control sampling sites impossible as the entire estuary was impacted. However, multiple evidence is available concerning the increase in sediment contamination and its impacts on benthic assemblages and effects on fish ecology due to the impact event (Gomes et al., 2017; Bernardino et al., 2019; Andrades et al., 2020). Qualitative fish sampling was conducted in the estuary in August, 2017 with the aid of a gillnet (5 cm internodes). Estuarine *Cathorops spixii* (n = 15), *Genidens genidens* (n = 18), *Eugerres brasilianus* (n = 18), *Diapterus rhombeus* (n = 9), and *Mugil* sp. (n = 11) specimens were captured, cryoanesthetized and stored at 4 °C until laboratory processing. The field sampling was conducted under SISBIO/ICMBio license number 64345-4. All fish sampled are demersal species with predominant benthic feeding habits (Andrades et al., 2020). After dissection, fish muscle and liver tissues were sampled and stored at −80 °C until analysis.

Surface sediment samples (0-5 cm) were randomly sampled at 17 stations along the estuary using a Van-Veen Grab and stored in previously decontaminated (30% HNO3) containers (Fig. 1). The location of samples are part of a long-term (4-year) monitoring of the estuary that include pre-impact and post-impact monitoring (Gomes et al., 2017; Bernardino et al., 2019); which provides a temporal evidence for the increased contamination of estuarine sediments associated with the tailings deposited after the disaster in 2015. Surface water physicochemical parameters, namely salinity, temperature, pH and total dissolved solids (TDS) were measured in situ using a HANNA HI9829 multi-parameter probe (Table 1).

**Figure 1.**
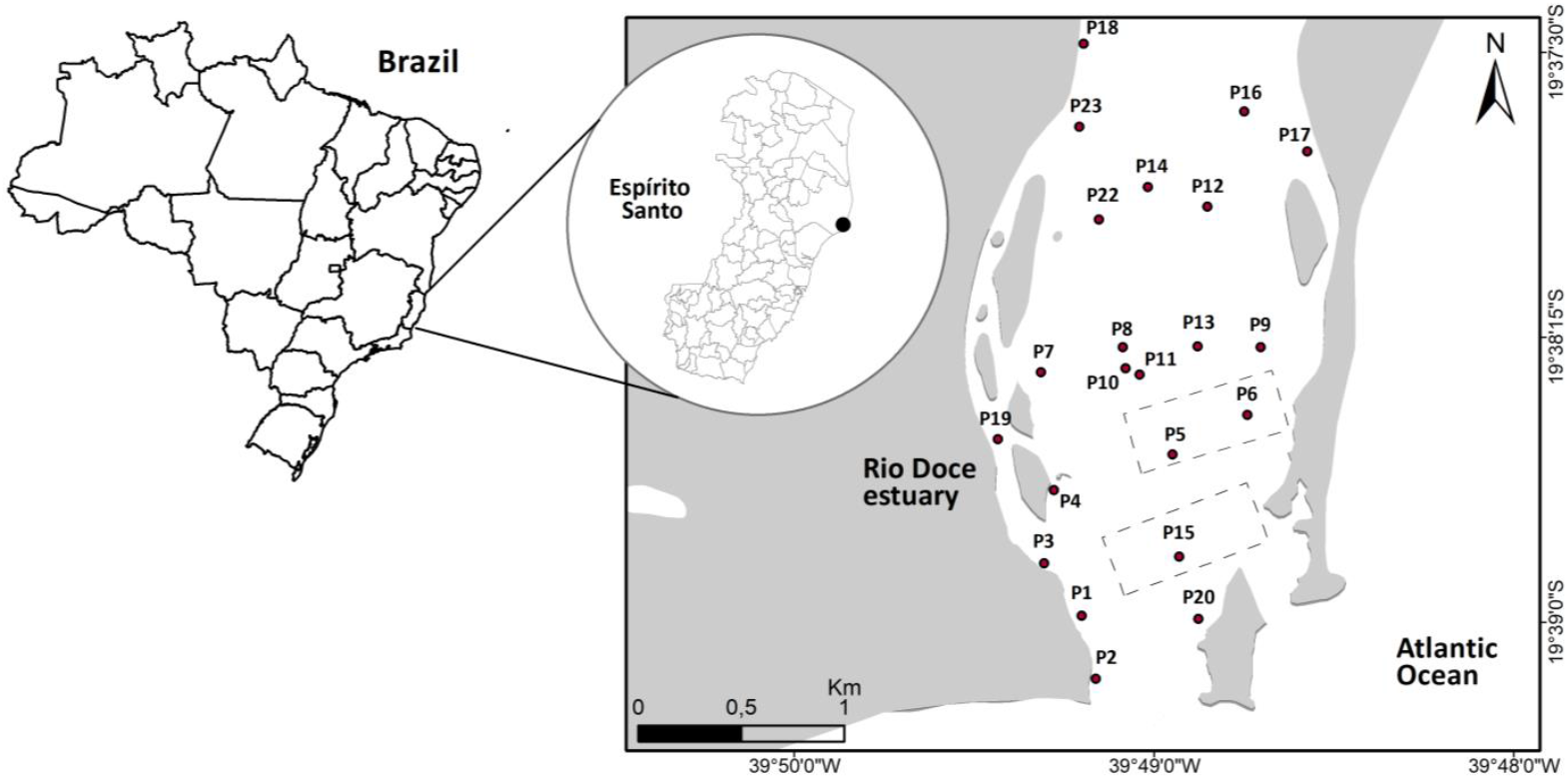
Map of the sediment sampling stations in the Rio Doce estuary, Brazil in August 2017 (black circles) and fish sampling areas (dotted rectangle).

**Table 1.** Environmental surface water parameters collected in August 2017 in the Rio Doce estuary. TDS - total dissolved solids.

|  | Salinity | Temperature (°C) | pH | TDS (ppm) |
| --- | --- | --- | --- | --- |
| Median | 0.27 ± 0.86 | 23.15 ± 0.75 | 7.35 ± 0.34 | 269 ± 779.06 |
| (min - max) | (0.10 - 3.7) | (22.3 - 24.9) | (6.5 - 8.1) | (99 - 3442) |

### Sediment analysis

Metals and metalloids in the sediment were determined by tri-acid digestion using HNO3, HF and H3BO3 in a microwave according to the EPA 3052 method (US EPA, 1996). The analysis included two-gram aliquots (wet weight) of the sediment. Digestion was performed using 9 mL of HNO3, 3 mL of HF (1 mol L^-1^) and 5 mL of H3BO3 (5%). Vessels containing the subsamples were shaken and heated at 110 °C for 4 hours. Subsequently, samples were diluted to 40 mL with deionized water. Finally, 0.1 mL aliquots were analyzed on an ICP-OES (Thermo Scientific - iCAP 6200). The analyses were performed in triplicate. To guarantee quality-control, standard solutions were prepared from dilution of certified standard solutions and certified reference materials used for comparison to measured and certified values (Table 2). Comparison of metal concentrations with sediment quality guidelines was realized (Table 3).

**Table 2.** Limits of detection quality assurance and quality control of total content determined by the USEPA 3052 method for the analyzed metals.

| Quality assurance | As | Cd | Cr | Cu | Pb | Zn |
| --- | --- | --- | --- | --- | --- | --- |
| Detection limit | 0.01 | 0.01 | 0.01 | 0.01 | 0.01 | 0.01 |
| Measured value | 1.068 | 0.9619 | 0.9608 | 1.012 | 0.9917 | 0.9989 |
| Certified value | 1 | 1 | 1 | 1 | 1 | 1 |
| Recovery (%) | 106.8 | 96.19 | 96.08 | 101.2 | 99.17 | 99.89 |

**Table 3.**
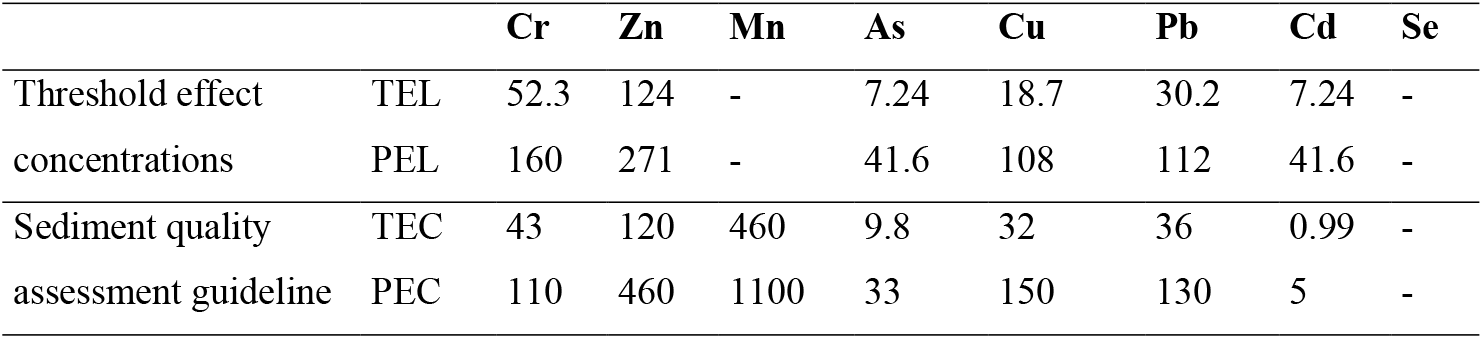
Sediment quality guidelines (SQGs). Concentrations expressed in mg kg-1. TEL - Threshold effect level, PEL - Probable effect level, TEC - Threshold effect concentrations, PEC - Probable effect concentrations.

|  |  | Cr | Zn | Mn | As | Cu | Pb | Cd | Se |
| --- | --- | --- | --- | --- | --- | --- | --- | --- | --- |
| Threshold effect | TEL | 52.3 | 124 | - | 7.24 | 18.7 | 30.2 | 7.24 | - |
| concentrations | PEL | 160 | 271 | - | 41.6 | 108 | 112 | 41.6 | - |
| Sediment quality | TEC | 43 | 120 | 460 | 9.8 | 32 | 36 | 0.99 | - |
| assessment guideline | PEC | 110 | 460 | 1100 | 33 | 150 | 130 | 5 | - |

### Metal and metalloid determinations in fish samples

Approximately 100 mg of wet samples (muscle and liver) were weighed in sterile polypropylene tubes, followed by the addition of 1.0 mL of bi-distilled HNO_3_. The blanks were prepared in triplicate, containing only 1.0 mL of bi-distilled HNO3. The DORM-4 (Dogfish muscle - National Research Council, Canada) Certified Reference Material (CRM) was used for quality control. The samples, blanks and CRM were left for approximately 12 hours overnight, then heated the following morning on a digester block for 4 hours at approximately 100 °C. The closed vessels were monitored hourly with manual pressure relief if necessary. After heating, the samples, CRM and blanks were left to cool at room temperature and made up to appropriate volumes with ultra-pure water (resistivity> 18MΩ). Element quantification was performed by ICP-MS using an ELAN DRC II ICP-MS (Perkin-Elmer Sciex, Norwalk, CT, USA). ^103^Rh was used as the internal standard at 20 μg L^-1^. The instrument limit of detection (LOD) and limit of quantification (LOQ) were estimated (Table 4).

**Table 4.** Instrumental limit of detection (LOD) and limit of quantification (LOQ) for the metals and metalloids determined in the present study (mg kg^-1^).

|  | Zn | Cu | Cd | Pb | Hg | As | Se | Cr | Mn |
| --- | --- | --- | --- | --- | --- | --- | --- | --- | --- |
| LOD | 0,0455 | 0,0060 | 0,0077 | 0,0038 | 0,0076 | 0,0283 | 0,0255 | 0,0730 | 0,0020 |
| LOQ | 0,1516 | 0,0200 | 0,0255 | 0,0126 | 0,0254 | 0,0943 | 0,0851 | 0,2433 | 0,0067 |

### Metallothionein (MT) extraction and determination

Samples were prepared according to the protocol proposed by Erk et al. (2002), with modifications. Briefly, muscle and liver samples (50 mg) were homogenized for 3 minutes in 300 μL of solution comprising 20 mmol L^-1^ Tris-HCl pH 8.6, phenylmethanesulphonyl fluoride 0.5 mmol L^-1^ as the antiproteolytic agent and β-mercaptoethanol 0.01% as the reducing agent. The samples were then centrifuged at 20,000 x g at 4 °C for 60 minutes. The resulting supernatants were separated from the pellets and placed in new microtubes. Proteins in the samples were denatured by heating the semi-purified supernatants for 10 minutes at 70 °C, followed by centrifugation for 30 minutes in the same conditions. Finally, the supernatants containing MT were transferred to new microtubes and frozen at −80 °C until analysis.

Metallothionein quantification via sulfhydryl content determination was performed by UV-Vis spectrophotometry through Ellman’s reaction (Ellman, 1959). Briefly, the samples were treated with a mixture of 1 mol L^-1^ HCl, 4 mol L^-1^ EDTA and 2 mol L^-1^ NaCl containing 5.5 dithio-bis (2-nitrobenzoic acid) buffered in 0.2 mol L^-1^ sodium phosphate pH 8.0. After incubation for 30 minutes, sample absorbances were determined at 412 nm on a UV-Vis spectrophotometer. Metallothionein concentrations were estimated using an analytical curve plotted with GSH as an external standard and transformed to metallothionein through the known stoichiometric relationship between metallothionein and reduced glutathione, of 1:20, as GSH contains 1 mol of cysteine per molecule and metallothionein, 20 moles.

### Reduced glutathione extraction and determination

The reduced glutathione analysis was carried out according to the protocol proposed by Beutler (1975), with modifications introduced by Wilhelm-Filho et al. (2005). Briefly, about 25 mg of each tissue were weighed and homogenized in 350 μL of 0.1 mol L^-1^ sodium phosphate buffer pH 6.5 containing 0.25 mol L^-1^ sucrose. The samples were then centrifuged at 11,000 x g for 30 minutes at 4 °C. The supernatants were transferred to microtubes and treated with 0.1 mol L^-1^ DTNB at pH 8.0 with a 1:1 ratio. After incubation for 15 minutes in the dark, sample absorbance was determined at 412 nm on a UV-Vis spectrophotometer. Reduced glutathione concentrations were estimated using an analytical curve plotted with GSH as an external standard (Monteiro et al., 2006).

### National and international references for fish consumption

Metal levels in fish muscle and liver tissues were compared to maximum permissible levels for consumption according to Brazilian (Brazilian Health Regulatory Agency - ANVISA, 1965) and international (Food and Agriculture Organization of the United Nations - FAO/WHO, 1997; American Food and Drug Administration - US FDA, 1993; Environmental Protection Agency - US EPA, 2007; British Ministry of Forestry, Agriculture and Fisheries (MAFF, 1995, and European Community legislation – EC, 2001) standards (Table 5).

**Table 5.**
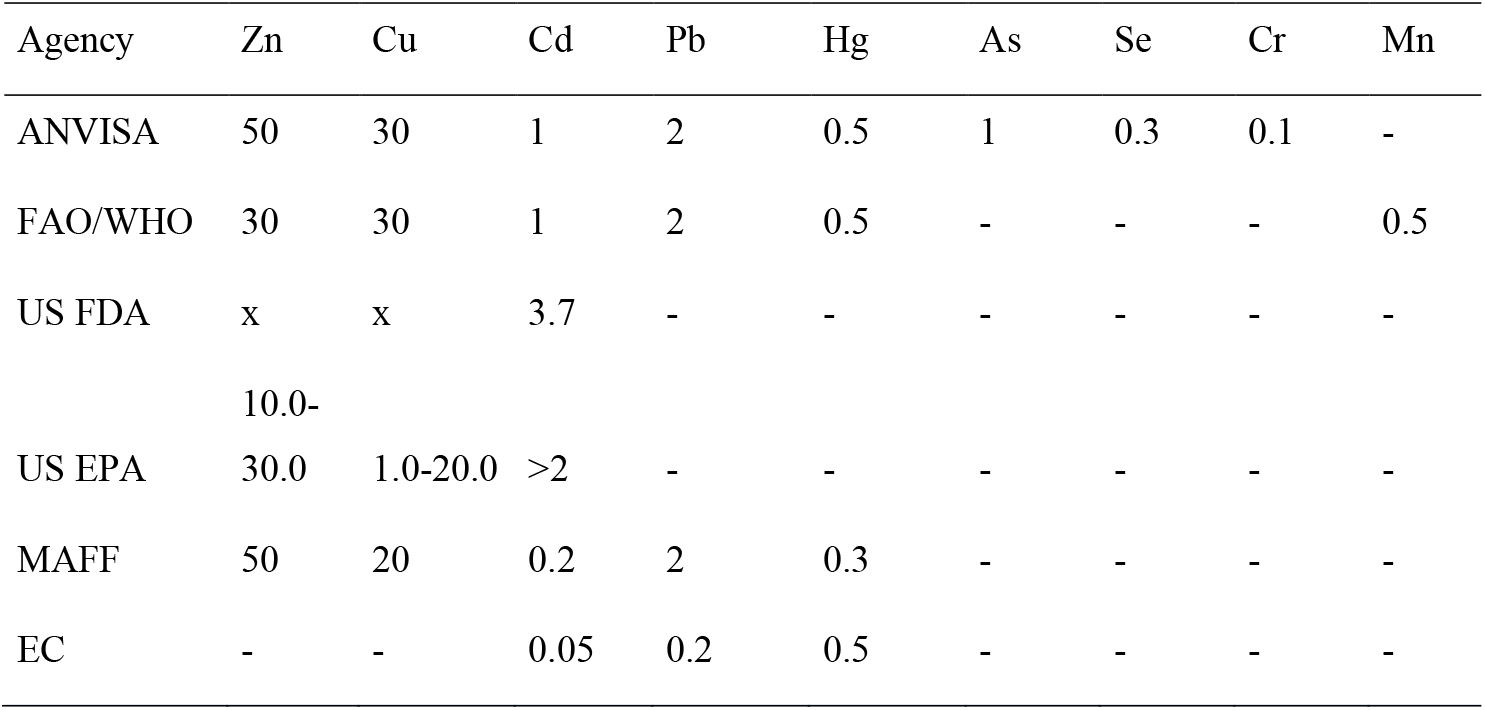
National and international maximum permissible levels (mg kg^-1^) for the ingestion of fish products in Brazil and worldwide.

### Statistical analyses

Data was expressed as means ± SD and descriptive statistical analyses performed using GraphPad 8.0.2 software (GraphPad Software, San Diego, California, USA). Before carrying out the statistical tests, data distribution was verified by the Shapiro-Wilk test. Since the data was normally distributed, parametric tests were applied. Pearson’s correlation test was used to verify the existence of significant correlations between metal and metalloid concentrations and metallothionein and reduced glutathione data. As no statistically significant differences were observed between fish size, weight, and sex, the groups were treated homogeneously without a weight/size stratification range or sex separation.

A Canonical Analysis of Principal Coordinates (CAP; Anderson & Willis, 2003) complemented by multidimensional scaling (Anderson, 2001; McArdle & Anderson, 2001; Oksanen et al., 2018) was performed in order to evaluate the metal contamination and stress protein expression. Data was square-root transformed prior to CAP. CAP was then used to identify the metal or group of metals that best explained the variation in stress protein expression among species, and to determine the protein that contributed most to the differences among samples. All statistical tests used an α = 0.05 significance level. Graphical and analytical processing was performed in Numbers (Apple Inc.) and R project (R Development Core Team 2005) using the ‘stats’ and ‘vegan’ packages (Oksanen et al., 2018).

## Results

### Trace-metal contamination and sediment quality assessments

The mean Cr, Zn, Mn, As, Cu, Pb, Cd and Se concentrations in sediments were often higher than the reference values (pre-impact) for the estuary (Gomes et al., 2017), indicating an accumulation of these elements since the initial impact in 2015 (Table 6). The concentrations reach 3.6 ± 1.4 mg kg^-1^ of Cd, 38.7 ± 14.5 mg kg^-1^ of Zn, 100.2 ± 47.5 mg kg^-1^ of Pb, and 47.4 ± 15.2 mg kg^-1^ of Cr. These concentrations are, respectively, 35,900%, 2,319%, 2,031%, and 1,217% higher than those corresponding to the average concentrations of the pre-impact. When compared to sedimentary quality guidelines, sedimentary Pb concentrations (min. 5.6 and max. 192.9 mg kg^-1^) were higher than the threshold effect level (TEL; 94% of stations), probable effect level (PEL; 47% of stations), threshold effect concentrations (TEC; 82% of stations) and probable effect concentrations (PEC; 23% of samples). Sedimentary Cr and As were higher than the TEL and PEC in 47-23 and 53-23% of the samples and 53% of the samples, respectively. Mn was higher than the TEC in 59% and Cd higher than both the TEC and PEC in 94 and 12% of the samples, respectively. The results indicate that Cr, Mn, As, Pb and Cd concentrations are likely to result in harmful effects on organisms. On a toxicity risk scale, PEC sediment concentrations were exceeded for Pb and Cd.

**Table 6.** Comparison between sediment quality guidelines (SQGs), Threshold effect level (TEL), Probable effect level (PEL), Threshold effect concentrations (TEC) and Probable effect concentrations (PEC) with the mean element values in sediments (in bold). Results are reported as mg kg^-1^. ^a^ Local reference values calculated from pre-impact assessment in the Rio Doce estuary by Gomes et al (2017). LOQ for Se and As = 0.01 mg kg^-1^.

| Estuary<br>samples N = 17 |  |  | Sediment quality guidelines (SQGs) |  |  |  |  |
| --- | --- | --- | --- | --- | --- | --- | --- |
|  |  |  | LRV<sup>a</sup> | Threshold effect<br>concentrations |  | Sediment quality<br>assessment guideline |  |
|  | min – max | Mean ± SD |  | TEL | PEL | TEC | PEC |
| Cr | 18-71.1 | **47.4**±15.2 | 3.6 | 52.3 | 160 | 43 | 110 |
| Zn | 18.2-78.1 | **38.7**±14.5 | 1.6 | 124 | 271 | 120 | 460 |
| Mn | 148.7-1002.7 | **540.8**±220 | 231 | - | - | 460 | 1100 |
| As | <LQ-28.8 | **7.8**±7.7 | 3.3 | 7.24 | 41.6 | 9.8 | 33 |
| Cu | 3-15.0 | **9.4**±4.0 | 1.3 | 18.7 | 108 | 32 | 150 |
| Pb | 5.6-192.9 | **100.2**±47.5 | 4.7 | 30.2 | 112 | 36 | 130 |
| Cd | 0.6-7.1 | **3.6**±1.4 | 0.01 | 7.24 | 41.6 | 0.99 | 5 |
| Se | <LQ-13.6 | **5.9**±4.3 | 1,0 | - | - | - | - |

### Biochemical biomarker responses and metal accumulation in fish samples

Metallothionein (MT) levels in the muscle tissue were similar to values in liver tissue in all sampled species (ANOVA, F= 2.816; p > 0.05). The reduced glutathione (GSH) concentrations were higher in liver in *C. spixii, G. genidens, D. rhombeus* and *Mugil* sp., and higher in muscle tissue in *E. brasilianus*. A significant difference in GSH levels was observed between liver and muscle tissues for *G. genidens* (ANOVA, F= 6.874; p<0.0001; Fig. 2).

**Figure 2.**
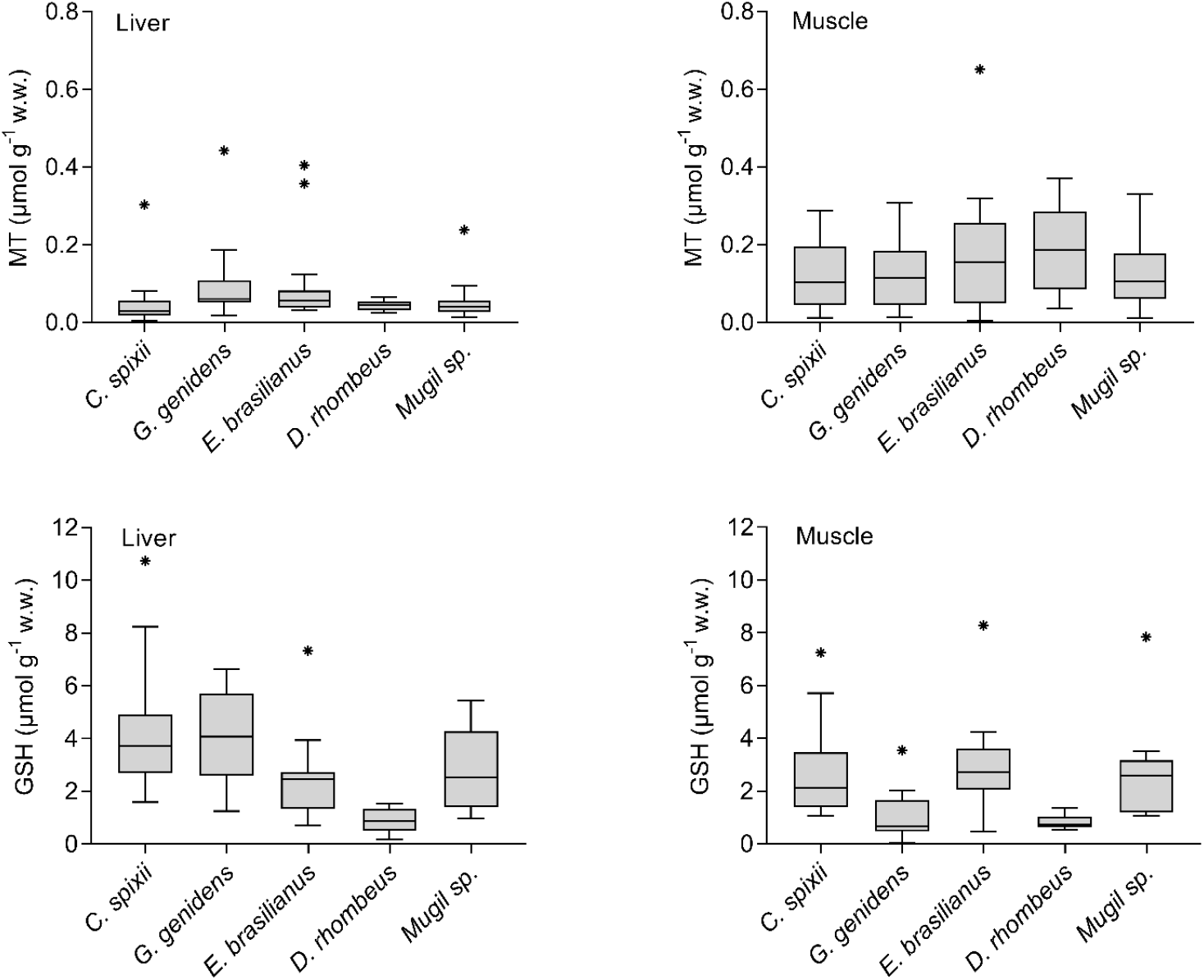
Total MT and GSH concentrations (μmol g^-1^ wet weight) in *C. spixii, G. genidens, E. brasilianus, D. rhombeus* and *Mugil* sp. liver and muscle tissues at the Rio Doce estuary. Box plots indicate minimum, maximum, median, quartiles, and outliers (asterisks).

Overall, higher metal concentrations in the liver compared to muscle tissues were observed. Tissue Cd concentrations were below the limit of quantification (LOQ = 0.0255 mg kg^-1^) in muscle in all fish species, while Pb displayed the same behavior only in *Mugil* sp. (Table 7). Zinc concentrations were significantly higher in the liver of all species (ANOVA, DF=99, F=22.9; p <0.0001), and Mn concentrations were higher in *E. brasilianus* liver (ANOVA, DF=19, F=30.51; p = 0.0309).

**Table 7.** Metal and metalloid concentrations in liver and muscle tissues of *C. spixii* (n = 15), *E. brasilianus* (n = 18), *Mugil* sp. (n = 11) and *D. rhombeus* (n = 9) sampled in the Rio Doce estuary in August 2017. ^a^ Indicates statistically significant differences between liver and muscle tissue (ANOVA, p <0.05). Limit of quantification (LOQ) in mg kg^-1^: Cd = 0.0255 and Pb = 0.0126.

| Species | Tissue | Zn | Cu | Cd | Pb | Hg | As | Se | Cr | Mn |
| --- | --- | --- | --- | --- | --- | --- | --- | --- | --- | --- |
| *Cathorops spixii* | Muscle | 8.21 ± 12.60<br>(3.58 - 46.64) | 0.3 ± 0.32<br>(0.14 - 1.27) | <LOQ<br><LOQ | 0.03 ± 0.02<br>(0.02 - 0.07) | 0.22 ± 0.02<br>(0.02 - 0.07) | 7.46 ± 10.46<br>(0.17 - 39.31) | 0.53 ± 0.2<br>(0.29 - 0.85) | 0.38 ± 0.14<br>(0.25 - 0.66) | 0.48 ± 0.97<br>(0.11 - 3.16) |
|  | Liver | 94.61 ± 203.05<sup>a</sup><br>(18.5 - 693.8) | 15.13 ± 37.87<br>(0.66 - 129.1) | 0.26 ± 0.17<br>(0.16 - 0.76) | 0.18 ± 0.11<br>(0.06 - 0.47) | 0.45 ± 0.52<br>(0.03 - 1.81) | 0.31 ± 1.24<br>(0.13 - 3.64) | 4.13 ± 3.42<br>(0.34 - 11.08) | 0.4 ± 0.11<br>(0.31 - 0.70) | 3.27 ± 1.84<br>(0.17 - 5.86) |
| *Genidens genidens* | Muscle | 14 ± 5.74<br>(6.61 - 95.41) | 0.29 ± 0.13<br>(0.11 - 0.61) | <LOQ<br><LOQ | 0.13 ± 0.07<br>(0.02 - 0.19) | 0.26 ± 0.15<br>(0.11 - 0.61) | 0.23 ± 4.12<br>(0.10 - 15.51) | 0.38 ± 0.08<br>(0.32 - 0.61) | 0.34 ± 0.11<br>(0.25 - 0.58) | 0.43 ± 0.19<br>(0.14 - 0.94) |
|  | Liver | 571.43 ± 320.2<sup>a</sup><br>(126.0 - 1310.9) | 8.94 ± 9.48<br>(1.89 - 46.2) | 0.41 ± 0.49<br>(0.03 - 2.36) | 0.29 ± 0.4<br>(0.03 - 1.91) | 0.75 ± 0.66<br>(0.14 - 3.02) | 0.21 ± 0.35<br>(0.12 - 1.35) | 3.91 ± 1.6<br>(2.37 - 9.75) | 0.41 ± 0.17<br>(0.30 - 0.62) | 1.49 ± 1.87<br>(0.89 - 9.00) |
| *Eugerres brasiliensis* | Muscle | 3.22 ± 4.61<br>(1.75 - 22.97) | 0.32 ± 0.40<br>(0.13 - 1.71) | <LOQ<br><LOQ | 0.055 ± 0.081<br>(0.023 - 0.23) | 0.19 ± 0.07<br>(0.07 - 0.30) | 0.20 ± 0.08<br>(0.11 - 0.331) | 0.69 ± 0.34<br>(0.33 - 1.90) | 0.39 ± 0.09<br>(0.26 - 0.51) | 0.35 ± 0.91<br>(0.08 - 4.22) |
|  | Liver | 27.27 ± 25.7<sup>a</sup><br>(3.41 - 119.56) | 2.36 ± 0.75<br>(0.32 - 3.66) | 0.19 ± 0.25<br>(0.04 - 0.96) | 0.05 ± 0.04<br>(0.02 - 0.14) | 0.11 ± 0.02<br>(0.06 - 0.15) | 0.51 ± 0.23<br>(0.09 - 0.91) | 2.14 ± 0.84<br>(0.58 - 3.86) | 0.42 ± 0.11<br>(0.25 - 0.67) | 7.20 ± 4.10<sup>a</sup><br>(0.28 - 17.25) |
| *Diapterus rhombeus* | Muscle | 3.81 ± 1.34<br>(2.5 - 7.1) | 0.16 ± 0.03<br>(0.14 - 0.22) | <LOQ<br><LOQ | 0.02 ± 0.01<br>(0.02 - 0.03) | 0.08 ± 0.05<br>(0.05 - 0.23) | 0.52 ± 0.56<br>(0.15 - 1.73) | 0.78 ± 0.23<br>(0.45 - 1.11) | 0.21 ± 0.04<br>(0.17 - 0.27) | 0.30 ± 0.53<br>(0.10 - 1.67) |
|  | Liver | 32.9 ± 28.68<sup>a</sup><br>(1.54 - 99.79) | 1.67 ± 0.82<br>(0.16 - 3.04) | 0.05 ± 0.26<br>(0.04 - 0.81) | 0.06 ± 0.02<br>(0.03 - 0.08) | 0.1 ± 0.03<br>(0.06 - 0.16) | 0.97 ± 0.62<br>(0.26 - 2.37) | 1.26 ± 0.79<br>(0.17 - 2.96) | 0.34 ± 0.11<br>(0.14 - 0.58) | 9.01 ± 8.29<br>(0.53 - 31.90) |
| *Mugil* sp. | Muscle | 3.21 ± 2.74<br>(1.81 - 12.18) | 0.30 ± 0.28<br>(0.15 - 0.73) | <LOQ<br><LOQ | <LOQ<br><LOQ | 0.07 ± 0.06<br>(0.05 - 0.24) | 0.25 ± 0.53<br>(0.17 - 2.02) | 0.50 ± 0.21<br>(0.17 - 0.97) | 0.32 ± 0.06<br>(0.27 - 0.42) | 0.20 ± 0.18<br>(0.07 - 0.57) |
|  | Liver | 71.71 ± 370.92<sup>a</sup><br>(23.8 - 1351.0) | 47.3 ± 150.6<br>(2.83 - 539.9) | 0.11 ± 0.35<br>(0.03 - 1.17) | 0.04 ± 0.3<br>(0.02 - 0.89) | 0.16 ± 0.4<br>(0.03 - 1.23) | 1.73 ± 1.99<br>(0.24 - 7.34) | 5.81 ± 9.33<br>(1.16 - 33.38) | 0.46 ± 0.17<br>(0.30 - 0.75) | 1.85 ± 2.14<br>(0.93 - 8.78) |

The CAP analysis indicated a significant association between trace metals in fish tissues and the expression of stress proteins (muscle F = 2.68, p = 0.016, liver F = 3.94, p = 0.003; Table 8, Fig. 3). In the liver, GSH expression was positively correlated to Zn and Hg concentrations (Zn F = 12.44, p = 0.003, Hg F = 12.42, p = 0.002; Table 8), mainly for *C. spixii* and *G. genidens* (Fig. 3). In muscle tissues, Cu and Cr displayed a higher contribution for GSH expression (Cu F = 7.12, p = 0.012, Cr F = 5.11, p = 0.028; Table 8), mainly for *C. spixii* and *E. brasilianus*. In general, differences in protein expression were the highest for *D. rhombeus* and the lowest for *Mugil* sp.

**Figure 3.**
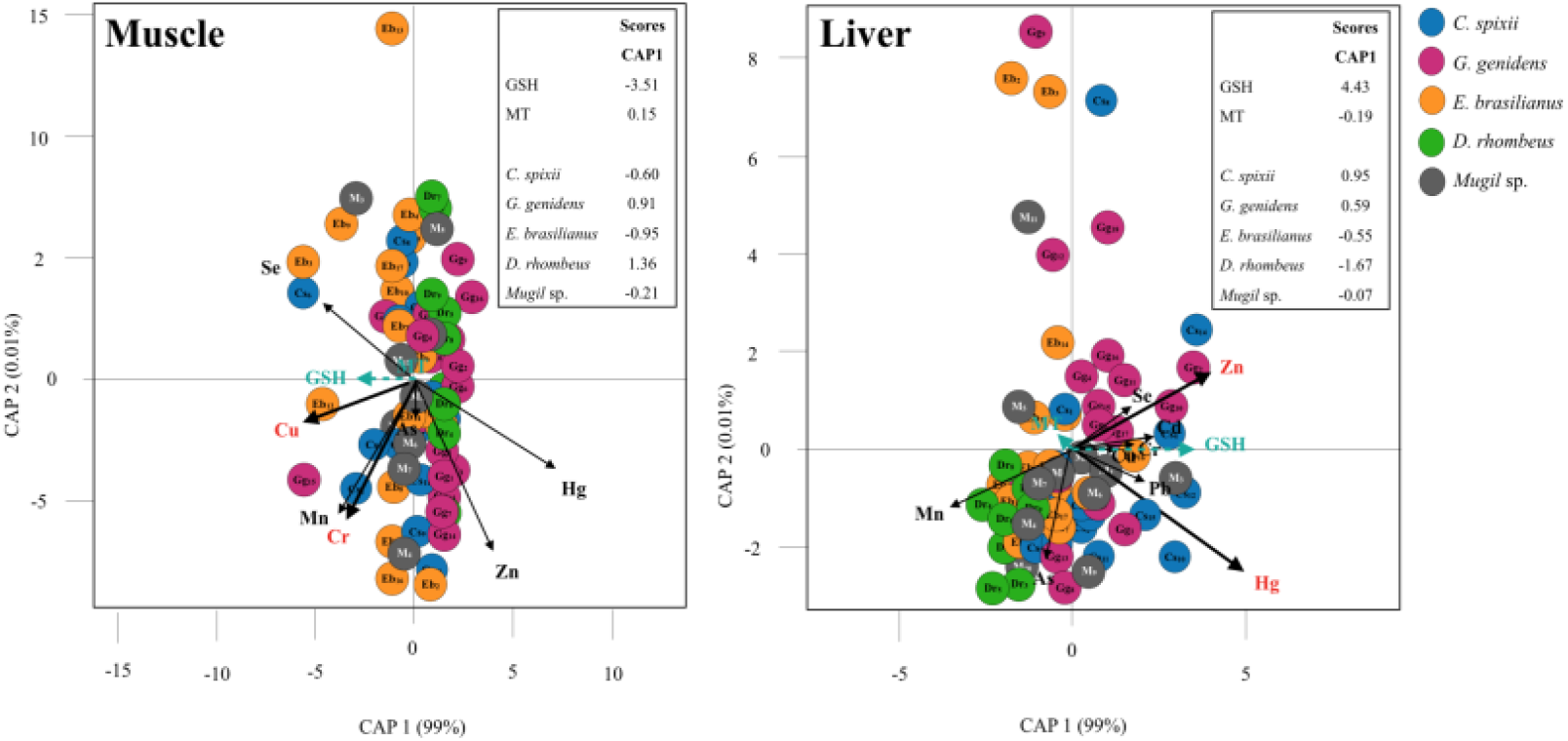
Canonical analysis of principal coordinates (CAP) indicating differences in expression of antioxidant biomarkers (GSH and MT) and the contribution of metal contamination (Zn, Cu, Cd, Pb, Hg, As, Se, Cr, Mn) in estuarine fishes (muscle and liver tissue). Vectors are based on Spearman correlation values > 0.5 (p < 0.5) for metals and scores for protein concentration and species (mean score among sampled). The proportion of data explained by axis 1 and 2 are in parenthesis.

**Table 8.** Results of the canonical analysis of principal coordinates (CAP) to evaluate the contribution of metal contamination (Zn, Cu, Cd, Pb, Hg, As, Se, Cr, Mn) and the variations in the expression of antioxidant biomarkers (GSH and MT) in estuarine fishes (muscle and liver tissue). Spearman correlation values for each metal are described for CAP axis 1-2. Note: proportion of variability explained by CAP axes are between parentheses ‘()’, Fisher test statistic, significant results (p < 0.05) are in bold.

|  | Muscle |  |  |  | Liver |  |  |  |
| --- | --- | --- | --- | --- | --- | --- | --- | --- |
| | F = 2.68, $p = 0.016$ | | | | F = 3.94, $p = 0.003$ | | | |
| | CAP 1<br>(0.99%) | CAP 2<br>(0.01%) | F | $p$ | CAP 1<br>(99%) | CAP 2<br>(0.01%) | F | $p$ |
| Zn | 0.28 | -0.69 | 1.61 | 0.200 | 0.59 | 0.22 | 12.44 | **0.003** |
| Cu | -0.52 | -0.14 | 7.12 | **0.012** | 0.18 | -0.002 | 1.61 | 0.203 |
| Cd | - | - | - | - | 0.33 | 0.04 | 0.29 | 0.613 |
| Pb | - | - | - | - | 0.31 | -0.09 | 2.76 | 0.097 |
| Hg | 0.58 | -0.40 | 3.03 | 0.084 | 0.74 | -0.37 | 12.42 | **0.002** |
| As | -0.29 | -0.12 | 0.51 | 0.490 | -0.11 | -0.34 | 2.29 | 0.153 |
| Se | -0.49 | 0.31 | 0.48 | 0.503 | 0.24 | 0.12 | 0.22 | 0.631 |
| Cr | -0.43 | -0.58 | 5.11 | **0.028** | 0.21 | 0.02 | 0.14 | 0.690 |
| Mn | -0.50 | -0.56 | 0.93 | 0.334 | -0.54 | -0.18 | 3.32 | 0.076 |

### Fish trace-element concentrations and human health standards

The trace metal wet weight results were compared to the maximum residue level (MRL) in food established by international agencies to evaluate human exposure to metals through the consumption of fish muscle and liver. Fish tissue contamination was compared to the most restrictive guideline for each element. High concentrations of Zn, Cu, Cd, Pb, Hg, As, Se, Cr and Mn in fish liver of all species analyzed exceeded guidelines, except for Pb and Hg in *D. rhombeus* and *E. brasilianus* (Table 9).

**Table 9.** Percentage (and number/total samples analyzed) of samples that exceeded maximum permissible levels allowed by Brazilian and international guidelines.

| Species | Matrix | Zn | Cu | Cd | Pb | Hg | As | Se | Cr | Mn |
| --- | --- | --- | --- | --- | --- | --- | --- | --- | --- | --- |
| *C. spixii* | Muscle | 33% (5/15) | 13% (2/15) | 0% (0/15) | 0% (0/15) | 20% (3/15) | 73% (11/15) | 87% (13/15) | 80% (12/15) | 53% (8/15) |
|  | Liver | 100% (15/15) | 93% (14/15) | 93% (14/15) | 13% (2/15) | 73% (11/15) | 40% (6/15) | 100% (15/15) | 93% (14/15) | 93% (14/15) |
| *G. genidens* | Muscle | 67% (12/18) | 0% (0/18) | 0% (0/18) | 0% (0/18) | 39% (7/18) | 33% (6/18) | 100% (18/18) | 100% (18/18) | 33% (6/18) |
|  | Liver | 100% (18/18) | 100% (18/18) | 94% (17/18) | 72% (13/18) | 89% (16/18) | 11% (2/18) | 100% (18/18) | 89% (16/18) | 100% (18/18) |
| *E. brasiliannus* | Muscle | 6% (1/18) | 11% (2/18) | 6% (1/18) | 6% (1/18) | 6% (1/18) | 0% (0/18) | 100% (18/18) | 44% (8/18) | 17% (3/18) |
|  | Liver | 94% (17/18) | 94% (17/18) | 83% (15/18) | 0% (0/18) | 0% (0/18) | 0% (0/18) | 100% (18/18) | 78% (14/18) | 94% (17/18) |
| *D. rhombeus* | Muscle | 0% (0/9) | 0% (0/9) | 0% (0/9) | 0% (0/9) | 0% (0/9) | 22% (2/9) | 100% (9/9) | 100% (9/9) | 33% (3/9) |
|  | Liver | 89% (8/9) | 89% (8/9) | 44% (4/9) | 0% (0/9) | 0% (0/9) | 44% (4/9) | 89% (8/9) | 100% (9/9) | 100% (9/9) |
| *Mugil* sp. | Muscle | 9% (1/11) | 9% (1/11) | 0% (0/11) | 0% (0/11) | 0% (0/11) | 9% (1/11) | 82% (9/11) | 36% (4/11) | 18% (2/11) |
|  | Liver | 100% (11/11) | 100% (11/11) | 82% (9/11) | 18% (2/11) | 36% (4/11) | 73% (8/11) | 100% (11/11) | 73% (8/11) | 100% (11/11) |

Liver tissue concentrations of Zn, Se and Mn exceeded guidelines for most specimens (>89%). Concentrations of As exceeded the guidelines for *C. spixii* in 73% of the analyzed specimens and in *G. genidens* for 33% of the muscle tissue samples. Cr concentrations exceeded guidelines in the muscle tissue of all analyzed specimens. Se concentrations also exceeded guidelines, both in liver and muscle tissue, in all *G. genidens* and *E. brasilianus* specimens, in *C. spixii* and *Mugil* sp. liver tissues, and in *D. rhombeus* muscle. Pb and Hg concentrations did not exceed the guidelines in *D. rhombeus*, but did in *C. spixii* and *G. genidens* liver and *E. brasilianus* muscle.

## Discussion

This study revealed tissue bioaccumulation of metals and metalloids and positive correlations to the expression of biomarkers of oxidative stress in fish obtained from the Rio Doce estuary 1.7 years after the mine tailing spill. Our results support the hypothesis of fish metal contamination and sublethal effects in aquatic ecosystems in the Rio Doce estuary, which were markedly augmented after the arrival of mine tailings in 2015. We detected tissue contamination in bottom-dwelling fish that are typically consumed by local populations, with potential health implications to villagers that rely on fish protein for their subsistence. Although baseline levels of trace metals in fish are unavailable for the estuary, it is very unlikely that the fish sampled 1.7 years after this disaster survived the acute impacts from the tailing in 2015 and would therefore, exhibit a legacy contamination concerning metals from the estuary. Hence, our findings suggest a rapid transfer (< 2 years) of trace metals from contaminated sediments to the biota.

The released mine tailing was initially characterized by low metal concentrations and with non-hazardous residues (Almeida et al., 2018). However, the tailings deposited in estuarine soils had significant concentrations of Fe oxides with high adsorption potential of toxic metals that were scavenged during their downstream riverine transport along 600 Km until it reached the estuary (Queiroz et al., 2018). Queiroz et al. (2018) hypothesized that the trace metals bound to Fe oxy-hydroxides could become bioavailable upon Fe reduction in estuarine soils, leading to high ecological risks to the estuarine biota (Gabriel et al., 2020). The mobilization of trace metals from tailings have occurred in aquatic riverine ecosystems downstream of the ruptured mining dam (Weber et al., 2020). Our results support the relatively rapid metal bioaccumulation in liver and muscle tissues of fish in aquatic ecosystems, and support that biogeochemical conditions in estuarine sediments may promote bioavailability of metals bound to Fe from tailing in the sediments.

The physiological responses of the investigated regarding metallothionein and reduced glutathione concentrations suggests sublethal metal contamination biological effects. These biomarkers play important roles in metal toxicokinetics through metal sequestration in tissues and subsequent organism detoxification (Forman, Zhang & Rinna et al., 2009; Ruttkay-Nedecky et al., 2013). The statistical correlations observed herein indicate a temporal response to contamination, through bioaccumulation processes through exposure throughout their lives to the presence of the tailings in the estuary. This is a key advantage of the use of these biomarkers, as most contaminated areas lack baseline studies as in the studied estuary. Thus, the observed biomarker responses are an additional support for the indication of chronic trace metal contamination effects in this ecosystem (Gusso-Choueri et al., 2018; Bernardino et al. 2019).

The general trend of higher MT concentrations in fish muscle observed in the present study may be associated with metal overloading in the liver and other excretory organs (e.g. kidneys), where excess metals accumulate in muscle (Pacheco et al., 2017; Souza et al., 2018), suggesting elevated exposure to the assessed contaminants. However, further monitoring of biomarker expression in fish and trace metal concentrations in fish muscle are required to confirm this hypothesis.

GSH expression in liver in *C. spixii, E. brasilianus, Mugil* sp., and *D. rhombeus* was correlated with Cd, Hg, Cr, Zn, and Mn, suggesting that GSH expression is related to oxidative stress caused by metals (Di Giulio et al., 1995; Atli & Canli, 2008; Monteiro et al., 2008; Sharma & Langer, 2014). Although GSH levels may vary among fish species, the fish captured in the Rio Doce estuary exhibited higher GSH expression when compared to fish in non-contaminated freshwater ecosystems upstream (Weber et al., 2020). In addition, contaminated freshwater ecosystems upstream exhibited similar effects of fish biomarker response (Weber et al., 2020) and caused internal degeneration of tissue in fishes exposed to tailings in the Rio Doce (Macedo et al., 2020). The GSH expression in fish captured in the estuary suggests that the local estuarine ecosystem health has also been severely compromised by metal contamination, with possible sub-optimal conditions for the development of fish species (Andrades et al., 2020).

Tissue accumulation of Cr, Zn, Mn, Cu, Pb, Cd, and Se was higher in the liver, as expected, as this is the primary metal detoxification organ (Hauser-Davis, Campos & Ziolli, 2012). Tissues like the liver, with a higher lipid content, can alert about the current accumulation of metals since the metals can reach it very quickly through the bloodstream after absorption and, thus, these concentrations are proportional to those present in the environment (Dural, Göksu & Özak, 2007; Lima-Júnior et al., 2012, Bosco-Santos & Luiz-Silva, 2020). Increased concentrations of metals in muscle tissue were in bottom-dwelling species *C. spixii* and *G. genidens*, may suggest a saturation response for metal and metalloid contamination (Lu et al., 2018; Souza et al., 2018). Several trace metals with high concentrations in fish muscle including Cu, Zn, Cd, and Hg were also observed at high concentrations in the mine tailings deposited in the estuary (Gabriel et al., 2020). The transfer of bioavailable metals from contaminated sediments in coastal ecosystems has been widely reported (Zhu et al., 2015; Hauser-Davis et al., 2016; Gusso-Choueri et al., 2018, Mason et al., 2019) and support that tailings in the Rio Doce estuary were the source of the observed sublethal effects and statistical correlations observed between metal and protein concentrations in fish.

Metal concentrations varied between fish species and between liver and muscle tissues, probably as a result of varied physiological responses and exposure to contaminants (Shah & Altindağ, 2005). The fishes sampled in the Rio Doce estuary have a similar demersal behavior and feed on benthic invertebrates and other food sources, which suggests direct ingestion of contaminated sediments and other pollutants (Dantas et al., 2019; Andrades et al., 2020). In addition, the fish behavior may increase exposure to contaminants in sediments during active search for food on the bottom, leading resuspension of contaminants and their intake through gills (Cline et al., 1994; Bustamante et al., 2003). The demersal catfish species of this region deserve special attention as they displayed the highest metal concentration in both hepatic and muscle tissues and may increase human health risks when consumed (Yi, Yang & Zhang, 2011). Differences in metal concentrations among species can also be associated to age, differences in metabolism or the presence of migratory behavior (Rodrigues et al., 2010). Fish age, in particular, may also reflect exposure periods in the environment and consequently may also influence metal concentrations. Thus, these factors, although not studied in detail herein deserve future investigation as they could support restrictions on fish consumption on vulnerable populations affected by the disaster.

Our study revealed that chronic metal contamination in the Rio Doce estuary lead to metal bioaccumulation, statistically correlated to the expression of detoxification proteins in fish. Several fish species sampled in the estuary nearly two years after the initial deposition of Fe-rich mine tailings were contaminated by toxic trace metals; although the estuary had been under historical human stress before the disaster in 2015. The expression of detoxifying biomarkers indicate current exposure to trace metals in aquatic ecosystems. Based on high trace metal contents in fish muscle, the consumption of demersal fish species poses risks to human health and should be prohibited in this estuarine region. Consumption of fish liver, including from *Mugil* spp., is considered a delicacy in some traditional communities (Hauser-Davis et al., 2016), and could highly increase contamination effects through human consumption. In addition, estuarine health is probably compromised by chronic contamination, likely to be sub-optimal for fish development and fisheries production, both of which are important indirect effects neglected in management actions.

## Conclusions

Significant metal fish tissue contamination and detoxification and oxidative stress defenses in fish were observed in response to contamination of the Rio Doce estuary by mine tailings. High concentrations of toxic metals in the liver of the demersal species *G. genidens* and *C. spixii* and their respective protein syntheses correlations indicate chronic sublethal effects, while higher metallothionein levels in muscle tissues suggests metal overload in excretion organs. Metal concentrations in both liver and muscle tissue were above Brazilian and international guidelines for Maximum Residue Limits in foods for Cr, Zn, Mn, As, Cu, Pb and Cd, indicating potentially high human risks if consumed by communities near the impacted areas. Although our study evaluated these effects nearly 2 years after the disaster, these effects are likely to continue as long as the tailing is deposited in the estuarine ecosystem, which will also likely offer sub-optimal conditions for the development of fish species.

## Acknowledgements

The authors would like to thank the Fundação de Amparo a Pesquisa e Inovação do Espírito Santo (FAPES), Conselho Nacional de Pesquisa e Desenvolvimento (CNPq) and Coordenação de Aperfeiçoamento de Pessoal de Nível Superior (CAPES) for the financial support granted to the Soil Benthos Rio Doce Network Project (FAPES 77683544/17) and the doctoral scholarship the lead author. AFB was also supported by a CNPq PQ grant 301191/2017-8. The authors also thank all colleagues who participated in the sampling work.

